## Supplementary Data for "Structural Diversity of B-Cell Receptor Repertoires along the B-cell Differentiation Axis in Humans and Mice"

### Species CDR-H3 template usage

To identify portions of structural space that were never seen in the human or mouse data, we searched for CDR-H3 clusters that were never utilized in the human and mouse data, recording all CDR-H3 templates belonging to these clusters. The number of such templates was 109 (~4% of all FREAD templates). Eighty-eight of the 109 unused CDR-H3 templates derived from nanobodies, which constituted ~32% of all nanobody CDR-H3 loops in our FREAD library. A further six unused templates belonged to engineered human single heavy domain antibodies. The remaining 15 templates were from conventional antibodies.

### Patterns of CDR-H3 cluster usage

We investigated conservatism of Structural Stem cluster usage between naïve and antigen-experienced BCR repertoires. We defined Structural Stem conservatism as the number of shared clusters between Structural Stem clusters in B-cell types (Supplementary Figure 13). In the human data, ~98% of Structural Stem CDR-H3 clusters from naïve BCR repertoires were found in antigen-experienced BCR repertoires. An analogous pattern was seen with the mouse data. Approximately 99% of Structural Stem CDR-H3 clusters in naive BCR repertoires were found in plasma IGHM BCR repertoires. Our results demonstrates that the same CDR-H3 clusters are preferentially over-represented across different B-cell types, with the number of these over-represented CDR-H3 clusters diminishing to none along the B-cell development axis. This again reinforces our findings that usage of CDR-H3 clusters becomes increasing different in BCR repertoires along the B-cell differentiation axis as only a small number of new over-represented CDR-H3 clusters are shared between antigen-experienced BCR repertoires. These over-represented clusters can be a product of antigen-specific clonally expanded B-cells.
