## Supplementary Figures for "Structural Diversity of B-Cell Receptor Repertoires along the B-cell Differentiation Axis in Humans and Mice"

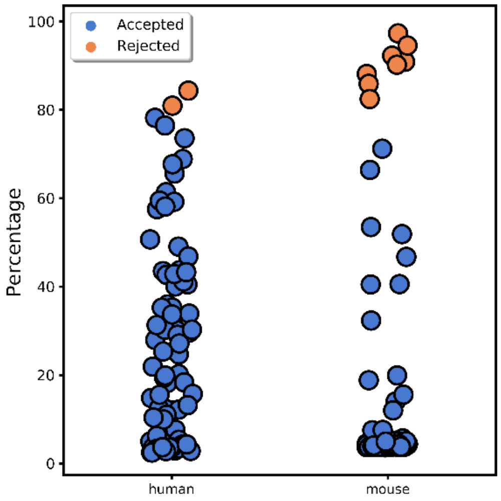


Supplementary Figure 1. **Percentage of sequences contained within the two most redundant CDR-H3 clusters for each BCR repertoire in the human and mouse data.** Each circle represents the percentage of sequences present in the two most redundant CDR-H3 clusters in a BCR repertoire. If this percentage exceeds 80% of the total number of that BCR repertoire's sequences (orange circle), the BCR repertoire was not included in the subsequent structural analysis. Different thresholds were checked, lower percentage cut-offs produced the same qualitative results in our structural analysis. Hence, 80% cut-off was selected to retain as many BCR repertories as possible.

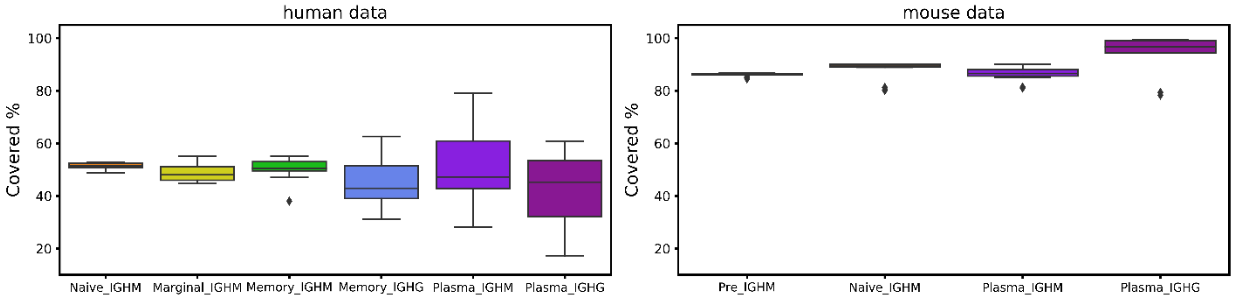


Supplementary Figure 2. **CDR-H3 structural coverage in human (A) and mouse (B) BCR repertoires.** FREAD CDR-H3 modellability was calculated across BCR repertoires of different B-cell types. FREAD modellability is defined as the percentage of sequences with predicted CDR-H3 structures over the total number of sequences in a given BCR repertoire.

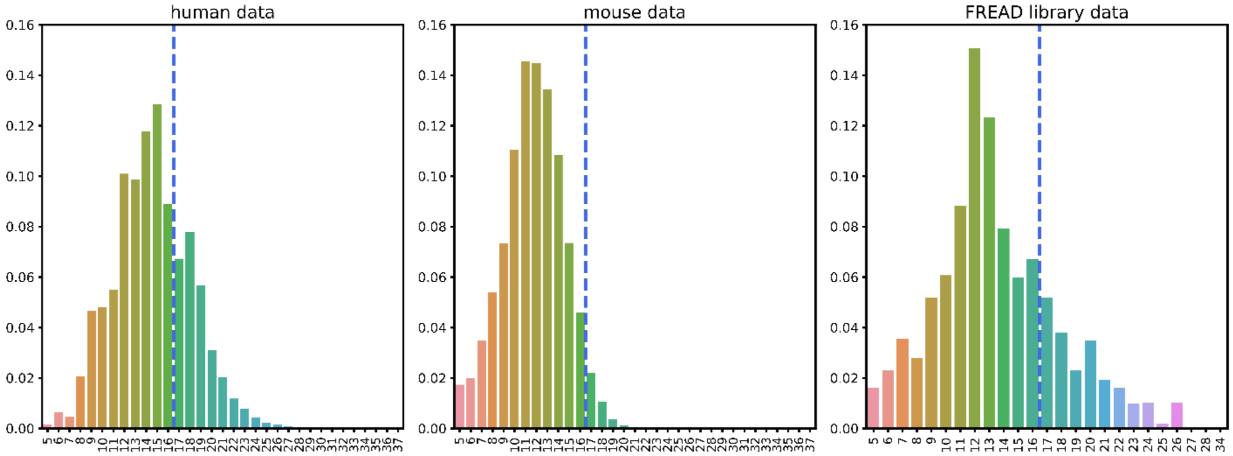


Supplementary Figure 3. **Normalized CDR-H3 length distribution in the human and mouse data, and our FREAD CDR-H3 template library**. Normalized CDR-H3 length distribution was calculated for all BCR repertoires in the human and mouse data, and all CDR-H3 templates that were in our FREAD library. Mouse CDR-H3s were on average shorter than human. The distribution of FREAD CDR-H3 lengths is different that found in the human and mouse data. The vertical blue line shows our CDR-H3 length cutoff (17 residues). Sequences and FREAD templates whose CDR-H3 lengths were longer than the cutoff were not considered in our analysis.

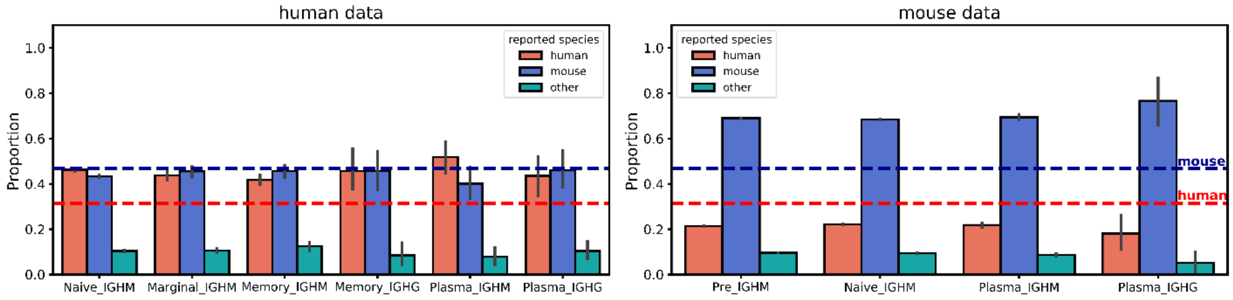


Supplementary Figure 4. **Reported species origin of CDR-H3 templates in the human and mouse data.** We calculated the proportion of CDR-H3 template reported species of origin for every BCR repertoire across different B-cell types. Reported species origin information was extracted from SAbDab (10). Orange bars show the proportion of human CDR-H3 templates, blue bars represent mouse CDR-H3 templates, while cyan bars depict the proportion of other than human or mouse CDR-H3 templates. The horizontal lines represent the expected outcome of uniform sampling of human (orange) and mouse (blue) CDR-H3 templates. Uniform sampling was calculated as the number of human or mouse CDR-H3 templates over the total number CDR-H3 templates found in our FREAD library.

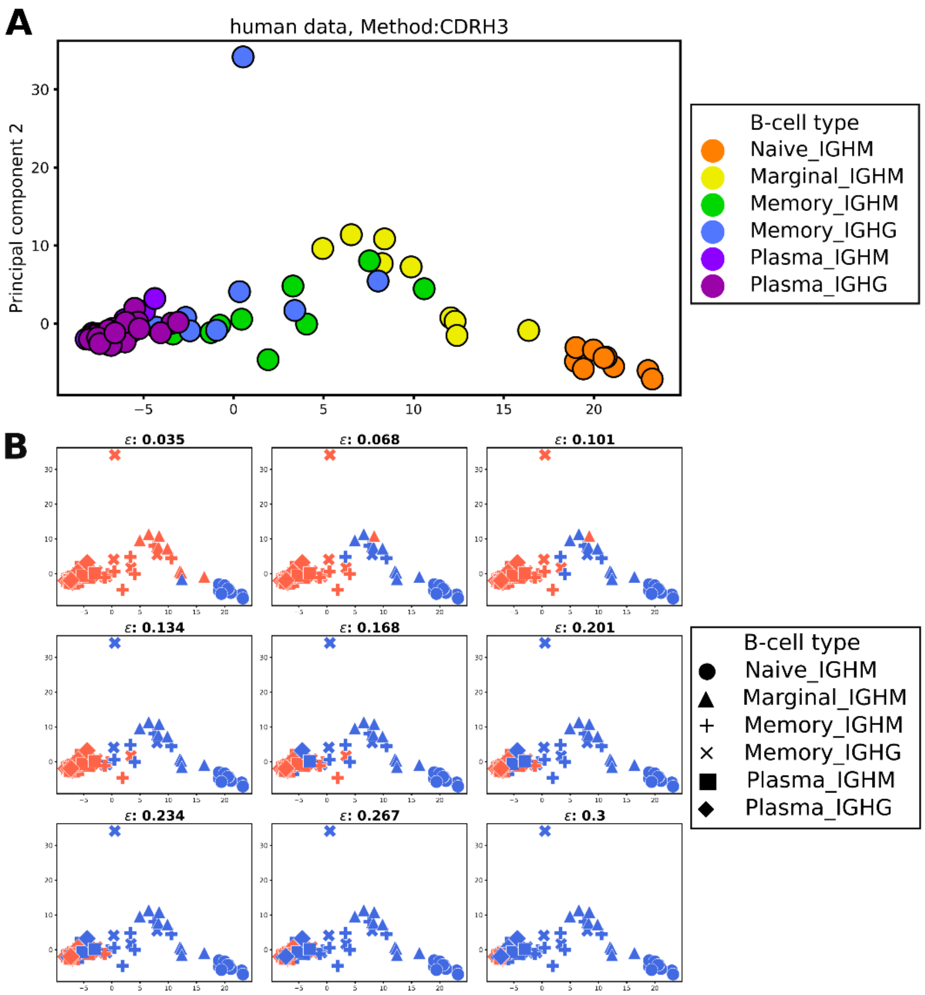


Supplementary Figure 5. **Analysis of densities of CDR-H3 cluster usage in the human data.** (**a**) PCA was performed on the human BCR repertoires (circles) with CDR-H3 cluster usages used as the features. The first two principal components were used to visualize any separation. Colours represent different B-cell types. (**b**) DBSCAN analysis with increasing maximum distance (ε) was employed to interrogate CDR-H3 cluster usage densities across human BCR repertoires. PCA analysis (as in **a**) was then used to visualize the DBSCAN clustering. The parameter ε was increased left-to-right, top-to-bottom. Marker shapes indicate different B-cell types; blue colour represents BCR repertoires that clustered with antigen-unexperienced BCR repertoires (naïve); orange colour shows DBSCAN-unclustered BCR repertoires at that ε value.

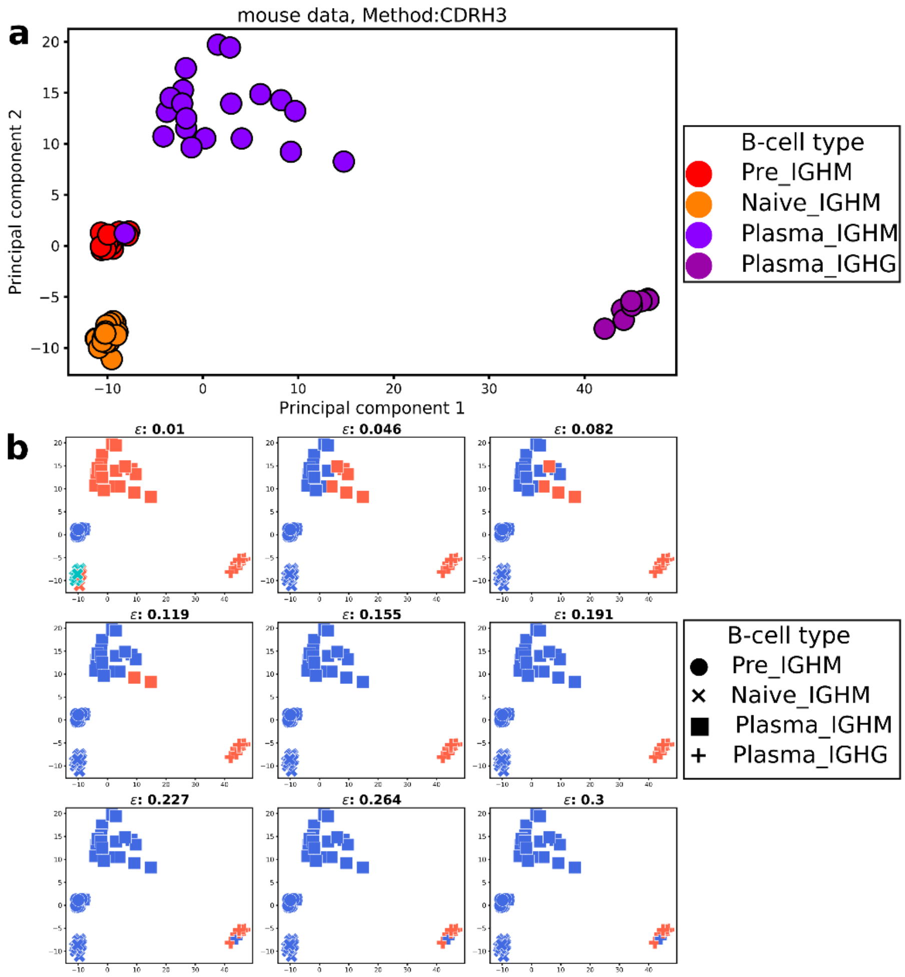


Supplementary Figure 6. **Analysis of densities of CDR-H3 cluster usage in the mouse data.** (**a**) PCA was performed on the mouse BCR repertoires (circles), with CDR-H3 cluster usages used as the features. The first two principal components were used to visualize any separation. Colours represent different B-cell types. (**b**) DBSCAN analysis with increasing maximum distance (ε) was employed to interrogate CDR-H3 usage densities across mouse BCR repertoires. PCA analysis (as in **a**) was then used to visualize the DBSCAN clustering. The parameter ε was increased left-to-right, top-to-bottom. Marker shapes indicate different B-cell types; cyan colour (in the top left subplot) represents naïve BCR repertoires, blue colour represents BCR repertoires that clustered with antigen-unexperienced BCR repertoires (pre and naïve); orange colour shows DBSCAN-unclustered BCR repertoires at that ε value.

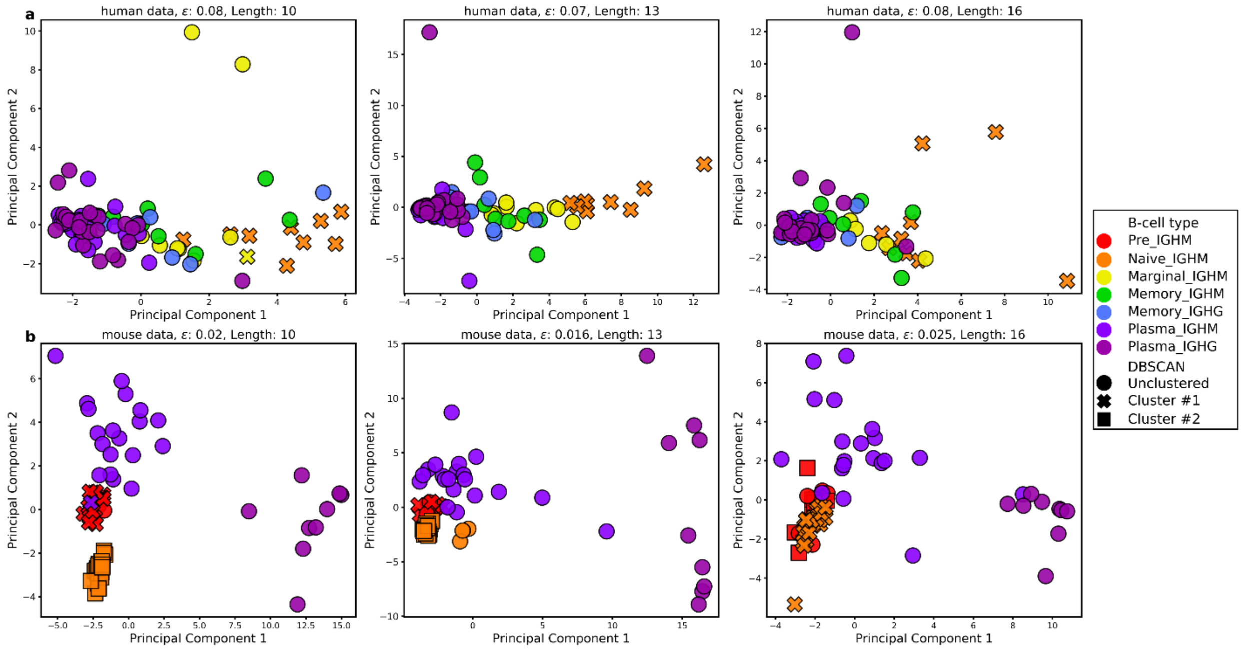


Supplementary Figure 7. **Structural interrogation of the human and mouse data with specific CDR-H3 lengths**. PCA was performed on **(a)** human and **(b)** mouse BCR repertoires, with CDR-H3 cluster usages selected as the features. The first two principal components were used to visualize any separation. Colours represent different B-cell types. DBSCAN was employed to quantify densities of CDR-H3 cluster usages across the repertoires. Marker shapes illustrate DBSCAN cluster information. Circle markers indicate DBSCAN unclustered BCR repertoires; other marker shapes show individual DBSCAN clusters. We considered a B-cell type separation if naïve and pre (antigen-unexperienced) BCR repertoires displayed the closest densities of CDR-H3 cluster usages at lower ε values in DBSCAN. In both **(a)** human and **(b)** mouse repertoires, antigen-unexperienced B-cell types cluster first regardless of CDR-H3 lengths. This confirms that BCR repertoires of different B-cell types have different patterns of CDR-H3 cluster usage.

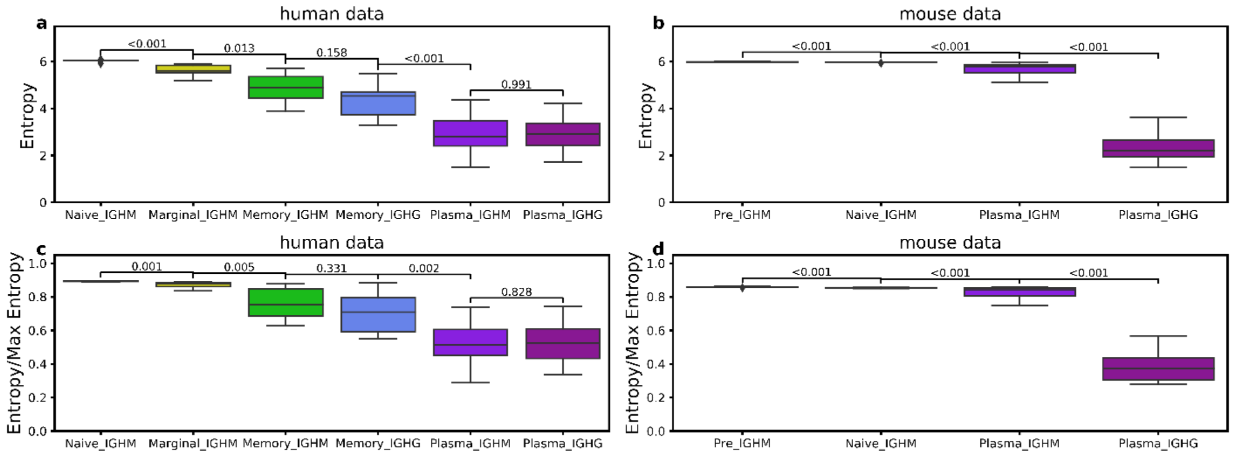


Supplementary figure 8. **CDR-H3 structural diversity in the human and mouse data.** Shannon entropy was calculated for CDR-H3 cluster usage in human (**a**) and mouse (**b**) data. To account for the varying numbers of CDR-H3 clusters across B-cell types, structural diversity of CDR-H3s was expressed as a proportion of entropy over theoretical maximum entropy in the human (**c**) and mouse (**d**) data. Theoretical maximum entropy was found for each BCR repertoire by using CDR-H3 structures represented in the given repertoire in equal proportions in entropy calculations. Higher values indicate higher diversities of CDR-H3 cluster usage. The Mann-Whitney U-test was used for statistical analysis and p-values are reported.

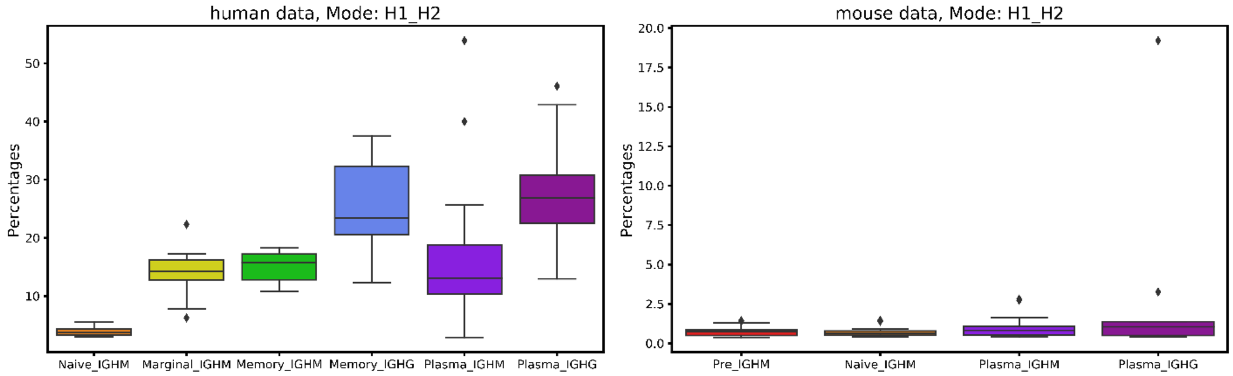


Supplementary Figure 9. **Canonical class divergence from parent germline class in the human and mouse data.** Canonical class divergence was defined as a mismatch in either the CDR-H1 or CDR-H2 canonical class from the germline canonical class. Percentages were calculated as the number of sequences with canonical class divergence over the total number of sequences in a given BCR repertoire. Colours represent different B-cell types.

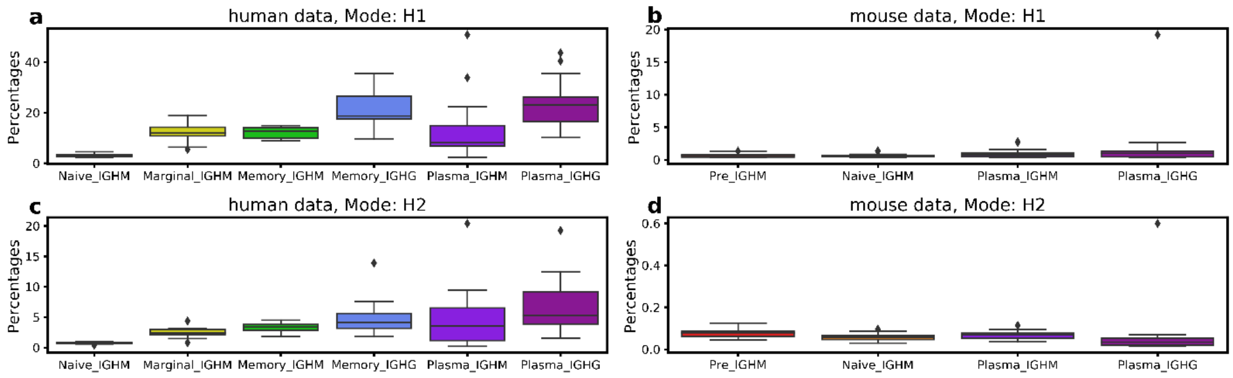


Supplementary Figure 10. **CDR-H1 and CDR-H2 canonical class divergence from parent germline class in the human and mouse data**. The top boxplots show canonical class divergence in CDR-H1 canonical class from the germline canonical class in the (**a**) human and (**b**) mouse data. Percentages were calculated as the number of sequences with canonical class divergence over the total number of sequences in a given BCR repertoire. The bottom boxplots show the same analysis on CDR-H2 canonical class in the (**c**) human and (**d**) mouse data. Colours represent different B-cell types.

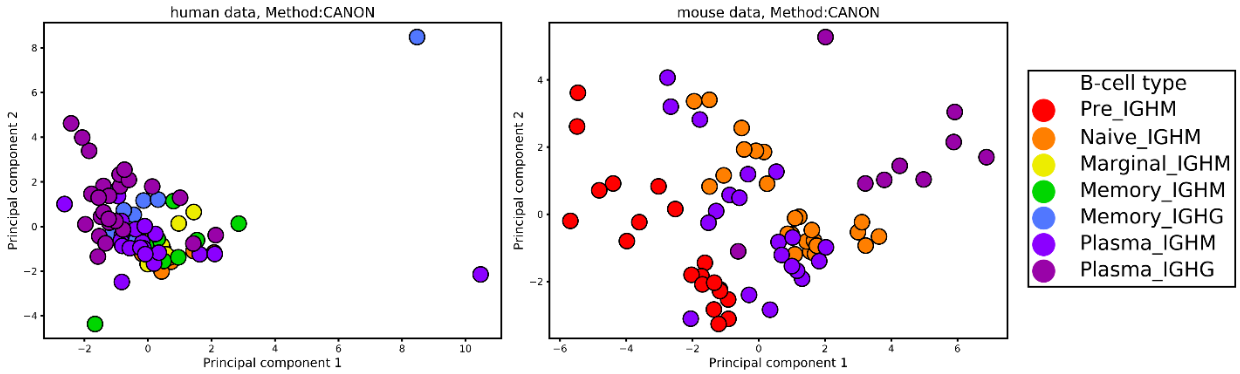


Supplementary Figure 11. **PCA on the human and mouse BCR repertoires.** Features included in the PCA were frequencies of CDR-H1 and CDR-H2 combinations in BCR repertoires. The first two principal components were used to visualize the separation of BCR repertoires. Colours represent different B-cell types.

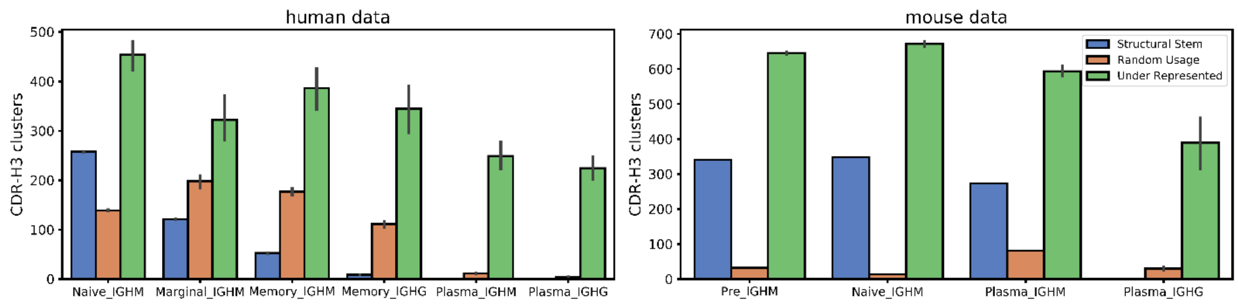


Supplementary Figure 12. **Number of CDR-H3 clusters based on their pattern of usage across different B-cell types in human and mouse data.** Structural Stems (blue bars) were defined as CDR-H3 clusters, which were over-represented across BCR repertoires of the same B-cell type. Under-Represented (green bars) were under-represented CDR-H3 clusters. CDR-H3 clusters, whose usages were not significantly different from random sampling, were termed Random-Usage (orange bars). The X-axis shows different B-cell types in the order of the B-cell maturation axis. The Y-axis shows the number of CDR-H3 clusters.

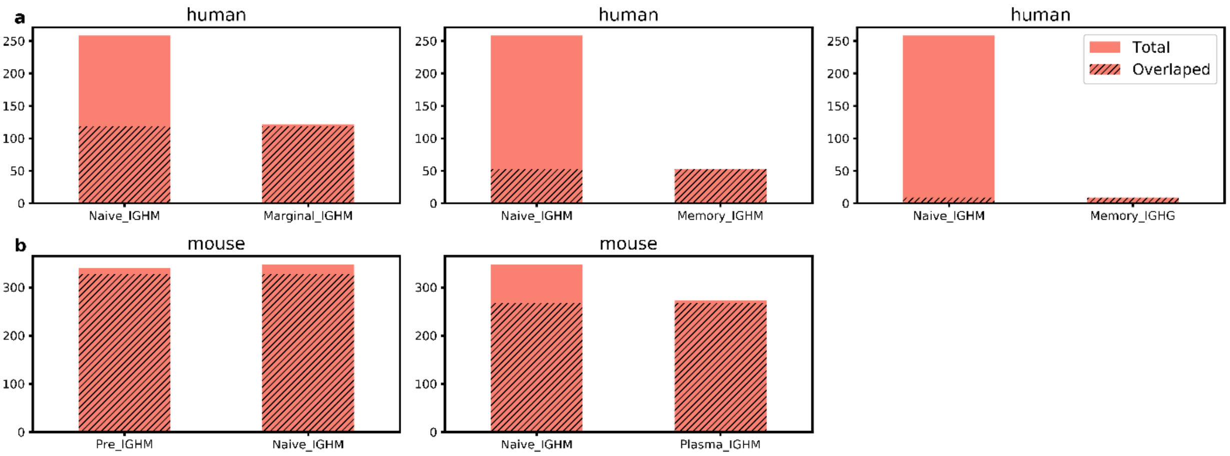


Supplementary Figure 13. **Overlap of Structural Stem CDR-H3 clusters between naïve and antigen experienced BCR repertoires in the human and mouse data**. Naïve and antigen experienced BCR repertoires were investigated for the Structural Stem overlap in the human (row **A**) and mouse (row **B**) data. The overlap was defined as a number of shared clusters between Structural Stem CDR-H3 clusters in BCR repertoires found in two different B-cell types. The X-axis shows the B-cell types. The Y-axis shows the total number of Structural Stem CDR-H3 clusters. The stripped pattern indicates the number of overlapped Structural Stem CDR-H3 clusters.
