## Supplementary Tables for "Structural Diversity of B-Cell Receptor Repertoires along the B-cell Differentiation Axis in Humans and Mice"

| Data | | |
|---|---|---|
| Human | RMSD | 2.5 Å |
| | Precision | 68.8% |
| Mouse | RMSD | 2.5 Å |
| | Precision | 68.4% |

Supplementary Table 1. **Estimated FREAD average RMSD and precision on the human and mouse data**. FREAD performance of CDR-H3 structure prediction was validated on the human and mouse data across three CDR-H3 length bins: 5 to 12, 13 and 14, and 15 and 16. For each length bin, ESS cutoffs were selected to achieve an average RMSD better than 3 Å or a coverage greater than 15%. The same ESS cutoffs were selected for both human and mouse data. Precision was defined as the percentage of FREAD predictions within 3 Å over the total number of predictions within the ESS cutoff.

| Data | Total sequences | CDR-H1 annotated | CDR-H2 annotated |
|---|---|---|---|
| Human | 5,712,939 | 5,598,599 (97.7%) | 5,425,279 (95.4%) |
| Mouse | 206,680,496 | 204,805,604 (99%) | 206,592,576 (~100%) |

Supplementary Table 2. **SCALOP annotation of Ig-seq data**. Annotation was performed on the human and mouse data. The human data contained 5.7 million sequences with CDR-H3 loop lengths of 16 amino acids or shorter. SCALOP predicted CDR-H1 loop shapes in 97.7% of sequences and CDR-H2 loop shapes in 95.4% in the human data. The total number of mouse sequences was ~207 million, of which 99% of CDR-H1 and ~100% of CDR-H2 loop shapes were annotated.

| Ig-seq data | B-cell type | Modelled | Total |
|---|---|---|---|
| human | Marginal_IGHM | 40542 | 79374 |
| human | Marginal_IGHM | 46155 | 96437 |
| human | Marginal_IGHM | 42817 | 85799 |
| human | Marginal_IGHM | 32810 | 71724 |
| human | Marginal_IGHM | 32903 | 73624 |
| human | Marginal_IGHM | 39059 | 85003 |
| human | Marginal_IGHM | 28438 | 59273 |
| human | Marginal_IGHM | 45568 | 88973 |
| human | Marginal_IGHM | 51067 | 92772 |
| human | Memory_IGHG | 26060 | 49960 |
| human | Memory_IGHG | 37809 | 71971 |
| human | Memory_IGHG | 19695 | 45813 |
| human | Memory_IGHG | 13814 | 33615 |
| human | Memory_IGHG | 17551 | 38234 |
| human | Memory_IGHG | 18077 | 56392 |
| human | Memory_IGHG | 22719 | 58293 |
| human | Memory_IGHG | 17350 | 55683 |
| human | Memory_IGHG | 29199 | 46650 |
| human | Memory_IGHM | 52804 | 100837 |
| human | Memory_IGHM | 34668 | 91278 |
| human | Memory_IGHM | 41848 | 84393 |
| human | Memory_IGHM | 40968 | 81141 |
| human | Memory_IGHM | 52362 | 98528 |
| human | Memory_IGHM | 51176 | 92876 |
| human | Memory_IGHM | 44921 | 89271 |
| human | Memory_IGHM | 48654 | 89175 |
| human | Memory_IGHM | 44027 | 93671 |
| human | Naive_IGHM | 32179 | 63320 |
| human | Naive_IGHM | 32287 | 66128 |
| human | Naive_IGHM | 36021 | 71747 |
| human | Naive_IGHM | 32499 | 63213 |
| human | Naive_IGHM | 33022 | 62914 |
| human | Naive_IGHM | 31464 | 60080 |
| human | Naive_IGHM | 30662 | 60381 |
| human | Naive_IGHM | 39334 | 75855 |
| human | Naive_IGHM | 35346 | 66803 |
| human | Plasma_IGHG | 26402 | 50206 |
| human | Plasma_IGHG | 24105 | 51982 |
| human | Plasma_IGHG | 29416 | 54956 |
| human | Plasma_IGHG | 17076 | 46453 |
| human | Plasma_IGHG | 21310 | 53057 |
| human | Plasma_IGHG | 10900 | 34201 |

| | | | |
|---|---|---|---|
| human | Plasma_IGHG | 21872 | 49349 |
| human | Plasma_IGHG | 18074 | 51268 |
| human | Plasma_IGHG | 27081 | 56119 |
| human | Plasma_IGHG | 15238 | 53452 |
| human | Plasma_IGHG | 24710 | 40603 |
| human | Plasma_IGHG | 19795 | 33585 |
| human | Plasma_IGHG | 28276 | 54691 |
| human | Plasma_IGHG | 18515 | 40902 |
| human | Plasma_IGHG | 26199 | 53475 |
| human | Plasma_IGHG | 25662 | 47180 |
| human | Plasma_IGHG | 12907 | 73711 |
| human | Plasma_IGHG | 15022 | 56705 |
| human | Plasma_IGHG | 13155 | 53597 |
| human | Plasma_IGHG | 34678 | 59596 |
| human | Plasma_IGHG | 28781 | 71801 |
| human | Plasma_IGHG | 14302 | 61565 |
| human | Plasma_IGHG | 16616 | 51762 |
| human | Plasma_IGHG | 40460 | 68094 |
| human | Plasma_IGHG | 35475 | 60455 |
| human | Plasma_IGHM | 39113 | 65219 |
| human | Plasma_IGHM | 60543 | 95026 |
| human | Plasma_IGHM | 47035 | 100136 |
| human | Plasma_IGHM | 37985 | 82823 |
| human | Plasma_IGHM | 31253 | 69234 |
| human | Plasma_IGHM | 92522 | 134775 |
| human | Plasma_IGHM | 45910 | 80793 |
| human | Plasma_IGHM | 45232 | 84968 |
| human | Plasma_IGHM | 46261 | 97241 |
| human | Plasma_IGHM | 15447 | 34514 |
| human | Plasma_IGHM | 40777 | 79151 |
| human | Plasma_IGHM | 31969 | 113387 |
| human | Plasma_IGHM | 27400 | 88127 |
| human | Plasma_IGHM | 34062 | 91438 |
| human | Plasma_IGHM | 41250 | 90472 |
| human | Plasma_IGHM | 70376 | 108498 |
| human | Plasma_IGHM | 20874 | 70800 |
| human | Plasma_IGHM | 26236 | 83401 |
| human | Plasma_IGHM | 87943 | 111339 |
| human | Plasma_IGHM | 66379 | 101631 |
| mouse | Naive_IGHM | 1342184 | 1664037 |
| mouse | Naive_IGHM | 1240472 | 1523085 |
| mouse | Naive_IGHM | 1426952 | 1775374 |
| mouse | Naive_IGHM | 4392652 | 4871630 |

| | | | |
|---|---|---|---|
| mouse | Naive_IGHM | 4061188 | 4514108 |
| mouse | Naive_IGHM | 13452930 | 15060651 |
| mouse | Naive_IGHM | 1583781 | 1967028 |
| mouse | Naive_IGHM | 4640306 | 5165988 |
| mouse | Naive_IGHM | 3427783 | 3805791 |
| mouse | Naive_IGHM | 4547582 | 5098906 |
| mouse | Naive_IGHM | 4497616 | 5013193 |
| mouse | Naive_IGHM | 4906734 | 5447184 |
| mouse | Naive_IGHM | 5166781 | 5740377 |
| mouse | Naive_IGHM | 4616391 | 5120337 |
| mouse | Naive_IGHM | 4378333 | 4860035 |
| mouse | Naive_IGHM | 2906410 | 3239617 |
| mouse | Naive_IGHM | 3678279 | 4076321 |
| mouse | Naive_IGHM | 2297687 | 2579810 |
| mouse | Naive_IGHM | 6414514 | 7127671 |
| mouse | Naive_IGHM | 3688098 | 4124532 |
| mouse | Naive_IGHM | 3109641 | 3458750 |
| mouse | Naive_IGHM | 6737178 | 7574357 |
| mouse | Naive_IGHM | 4137679 | 4585687 |
| mouse | Naive_IGHM | 6831077 | 7573020 |
| mouse | Plasma_IGHG | 395729 | 505410 |
| mouse | Plasma_IGHG | 190971 | 192559 |
| mouse | Plasma_IGHG | 672937 | 680672 |
| mouse | Plasma_IGHG | 254991 | 257230 |
| mouse | Plasma_IGHG | 214551 | 215655 |
| mouse | Plasma_IGHG | 214787 | 225339 |
| mouse | Plasma_IGHG | 276193 | 285690 |
| mouse | Plasma_IGHG | 85663 | 90681 |
| mouse | Plasma_IGHG | 354565 | 446868 |
| mouse | Plasma_IGHM | 1413089 | 1622047 |
| mouse | Plasma_IGHM | 1302080 | 1472598 |
| mouse | Plasma_IGHM | 7343414 | 8411665 |
| mouse | Plasma_IGHM | 1428102 | 1610945 |
| mouse | Plasma_IGHM | 1208881 | 1390785 |
| mouse | Plasma_IGHM | 2052537 | 2320385 |
| mouse | Plasma_IGHM | 1556711 | 1763522 |
| mouse | Plasma_IGHM | 2262469 | 2612130 |
| mouse | Plasma_IGHM | 1720837 | 1967491 |
| mouse | Plasma_IGHM | 2034987 | 2361737 |
| mouse | Plasma_IGHM | 2583351 | 2994422 |
| mouse | Plasma_IGHM | 1159962 | 1331034 |
| mouse | Plasma_IGHM | 1470144 | 1630570 |
| mouse | Plasma_IGHM | 1091139 | 1279155 |

| | | | |
|---|---|---|---|
| mouse | Plasma_IGHM | 476773 | 558585 |
| mouse | Plasma_IGHM | 1357383 | 1582355 |
| mouse | Plasma_IGHM | 1155550 | 1301587 |
| mouse | Plasma_IGHM | 2681711 | 3116805 |
| mouse | Plasma_IGHM | 1204579 | 1384501 |
| mouse | Plasma_IGHM | 1739957 | 2028798 |
| mouse | Pre_IGHM | 2019126 | 2326588 |
| mouse | Pre_IGHM | 2355741 | 2719441 |
| mouse | Pre_IGHM | 1263471 | 1478508 |
| mouse | Pre_IGHM | 1973160 | 2276983 |
| mouse | Pre_IGHM | 1728568 | 2006581 |
| mouse | Pre_IGHM | 1533743 | 1777646 |
| mouse | Pre_IGHM | 1893928 | 2198055 |
| mouse | Pre_IGHM | 787166 | 906148 |
| mouse | Pre_IGHM | 2593013 | 2989135 |
| mouse | Pre_IGHM | 2219150 | 2566267 |
| mouse | Pre_IGHM | 2306467 | 2666874 |
| mouse | Pre_IGHM | 1463339 | 1695145 |
| mouse | Pre_IGHM | 1830457 | 2116217 |
| mouse | Pre_IGHM | 1369005 | 1616993 |
| mouse | Pre_IGHM | 2424570 | 2799215 |
| mouse | Pre_IGHM | 1900629 | 2200358 |
| mouse | Pre_IGHM | 1689100 | 1957440 |
| mouse | Pre_IGHM | 1506394 | 1770037 |
| mouse | Pre_IGHM | 3363456 | 3877874 |
| mouse | Pre_IGHM | 2702801 | 3126281 |

Supplementary Table 3. **BCR repertoires used in structural diversity analysis**. Ig-seq data columns states species origin of Ig-seq data; modelled columns shows the number of BCR repertoire sequences with predicted CDR-H3 structures; Total column shows the total number of sequences in a given BCR repertoire whose CDR-H3 lengths are between 5 and 16 amino acids.
