## Supplementary Material for "Structural Diversity of B-Cell Receptor Repertoires along the B-cell Differentiation Axis in Humans and Mice"

### Supplementary Methods

#### FREAD Performance Assessment

The method used by FREAD to map an input loop sequence onto a known loop structure relies on the compatibility of their anchor region orientation as well as a sequence similarity score (Environment-specific Substitution Score (ESS)). ESS scores are calculated based on observed amino acid substitution probabilities in a protein evolutionary environment (1).

CDR-H3s exhibit high sequence and length diversity in Ig-seq data (2). Our FREAD library contains CDR-H3 loop structures derived from the SAbDab (3). The sequence diversity of these CDR-H3 templates is not representative of Ig-seq data, since the most these were subjected to rational antibody engineering (4), and their length distribution is different from natural Ig-seq data (Supplementary Figure 3). Therefore, we expect relatively low ESS scores between an Ig-seq sequence and its best hit in the FREAD library.

To accurately estimate FREAD performance for CDR-H3 structure prediction on a given Ig-seq dataset, we used the following three-step method. First, we used FREAD to predict templates for all SAbDAb CDR-H3 sequences, retaining all suggested templates alongside their ESS scores ('all versus all'). Here, we allowed FREAD to suggest templates with identical CDR-H3 sequences, since these sequences can be observed in natural Ig-seq data (5). To evaluate the structural similarity between each FREAD template and the native loop, we measured backbone RMSD using the DTW algorithm (6). This yielded a distribution of ESS scores with accompanying probabilities of RMSD values.

Next, we generated the combined distribution of ESS scores across the top FREAD predictions in the human and mouse Ig-seq data (ESS_TOP). Since the mouse data contains ~66 times more sequences, it was subsampled to match the number of sequences found in the human data. To ensure that representative murine ESS scores were picked, this subsampling procedure was repeated 100 times and the average ESS scores were recorded.

Finally, we randomly picked 1000 FREAD predictions by ESS score (with replacement) from our all versus all assessment to match the ESS_TOP distribution. This was repeated 100 times and the average RMSD and precision scores were calculated across different loop length bins (5 to 12, 13 and 14, and 15 and 16) and ESS values. This generated the distribution of CDR-H3 length bins with accompanying average RMSD (bins_RMSD) and precision (bins_precision) values.

We chose ESS cutoffs to achieve an average RMSD better than 3 Å or at least 15% of coverage for every length bin in the human and mouse data. Supplementary Table 1 shows the average RMSD and precision we estimate FREAD will achieve on the Ig-seq data.
